# Experimental Test of the Contributions of Initial Variation and New Mutations to Adaptive Evolution in a Novel Environment

**DOI:** 10.1101/2022.05.31.494207

**Authors:** Minako Izutsu, Richard E. Lenski

## Abstract

Experimental evolution is an approach that allows researchers to study organisms as they evolve in controlled environments. Despite the growing popularity of this approach, there are conceptual gaps among projects that use different experimental designs. One such gap concerns the contributions to adaptation of genetic variation present at the start of an experiment and that of new mutations that arise during an experiment. The primary source of genetic variation has historically depended largely on the study organisms. In the long-term evolution experiment (LTEE) using *Escherichia coli*, for example, each population started from a single haploid cell, and therefore adaptation depended entirely on new mutations. Most other microbial evolution experiments have followed the same strategy. By contrast, evolution experiments using multicellular, sexually-reproducing organisms typically start with pre-existing variation that fuels the response to selection. New mutations may also come into play in later generations of these experiments, but it is generally difficult to quantify their contribution in these studies. Here, we performed an experiment using *E. coli* to compare the contributions of initial genetic variation and new mutations to adaptation in a new environment. Our experiment had four treatments that varied in their starting diversity, with 18 populations in each treatment. One treatment depended entirely on new mutations, while the other three began with mixtures of clones, whole-population samples, or mixtures of whole-population samples from the LTEE. By tracking genetic markers associated with particular founders in two of our treatments, we could document the impact of the initial variation during the early generations of our experiment. However, there were no differences in fitness among the treatments after 500 or 2000 generations in the new environment, despite the variation in fitness among the founders. These results indicate that new mutations quickly overcame, and eventually contributed more to adaptation, than did the initial variation. Our study thus shows that pre-existing genetic variation can have a strong impact on early evolution in a new environment, but new beneficial mutations may contribute more to later evolution and can even drive some initially beneficial variants to extinction.

## INTRODUCTION

Some basic evolutionary issues can lead to misunderstandings and confusion, even among experts. One such issue concerns the contributions of standing genetic variation and new mutations to the process of adaptation by natural selection in a new environment. In this context, standing genetic variation includes those alleles that existed in a population before it encountered selection in the new environment, whereas new mutations are those alleles that arose after that selection began. It is a vexing problem because all genetic variation starts as new mutations and later can become standing variation, but the timing is important for understanding both the dynamics of evolutionary change within any single lineage and the repeatability of evolutionary outcomes across multiple lineages. With respect to the repeatability of evolution, Stern (2013) proposed the new term “collateral evolution” in juxtaposition with the more familiar idea of “parallel evolution” to emphasize how these different sources of genetic variation could lead to repeatable outcomes. Collateral evolution occurs when repeatable phenotypic changes evolve from standing variation in a common ancestral gene pool (i.e., variation that is identical by descent), whereas parallel evolution occurs when the similar phenotypes originate from independent mutational events (i.e., new mutations).

There is no single “right” answer in terms of the relative importance of standing variation and new mutations because both can contribute sequentially, simultaneously, and even synergistically to the process of adaptation by natural selection. But the ways that we do science—both conceptually and empirically—often lead us to emphasize one or the other source of genetic variation. In the long-term evolution experiment (LTEE) using *E. coli*, for example, new mutations are emphasized because each replicate population was founded from a single haploid cell of the ancestral strain in order to ensure that any repeatable outcomes result from independent mutations and hence parallel, rather than collateral, evolution (Lenski et al., 1991; Tenaillon et al., 2016; Lenski, 2017a). Hence, there was no standing variation at the start of the LTEE, and all of the genetic variation was produced by new mutations after the experiment began. Much of the work in the field of experimental evolution now follows the same mutation-dependent strategy, including most studies that use microorganisms (Tenaillon et al., 2012; Johnson et al., 2021). However, that approach is generally not followed in evolution experiments that use multicellular, sexually-reproducing plants and animals (Scarcelli and Kover, 2009; Burke et al., 2010; Schulte et al., 2010), for two largely practical reasons. First, quantitative genetics theory, which was developed for sexual plants and animals, presumes within-population genetic variation (Roff, 1997). That theory has guided artificial selection experiments to produce organisms with beneficial phenotypes for agricultural and other human applications (Hill and Caballero, 1992; Wright et al., 2005; Akey et al., 2010). By starting experiments with large, outbred populations that harbor abundant standing genetic variation, plant and animal breeders can improve traits more quickly than with small, inbred populations that lack diversity. Thus, most quantitative-genetic theories and applications presume that adaptation relies on standing variation, whereas the input from new mutations is typically ignored or abstracted (Roff, 1997). Second, the long generation times and small population sizes of larger organisms make evolution experiments that depend on new mutations (e.g., using near-isogenic inbred lines) impractical in most cases. Some studies using isogenic *Drosophila* populations failed to observe repeatable evolutionary changes (Harshman and Hoffmann, 2000), and relying on new mutations for adaptation in populations with long generation times requires experiments that are longer than most researchers are willing to perform (Izutsu et al., 2012). Therefore, researchers studying animals and plants usually start with outbred populations that harbor abundant genetic variation, and thus they have largely observed collateral evolution with respect to the repeatability of changes across replicate populations (Rose, 1984; Hoffmann et al., 2003; Mery and Kawecki, 2002; Barrett et al., 2008; Scarcelli and Kover, 2009; Burke et al., 2010; Schulte et al., 2010; Zhou et al., 2011; Graves et al., 2017).

In this study, we directly compare the rates of adaptation based on standing genetic variation versus new mutations, in order to fill the gap among studies using different model systems. To that end, we used various sets of bacteria from the LTEE as founders, and we then propagated them in a novel environment in which D-serine replaced glucose as the limiting resource. We had 18 populations in each of four treatments (Figure 1). In the Single-Clone (SC) treatment, each population started from a single clone sampled from one of six LTEE populations. In the Single-Population (SP) treatment, each population started from an entire LTEE population and all of the genetic variation present in that population. In the Mixed-Clones (MC) treatment, each population started as an admixture of the six SC founding clones. Finally, in the Mixed-Populations (MP) treatment, each population started as an admixture of the six SP founding populations. Thus, the SC populations did not have any initial within-population genetic variation, and therefore they relied entirely on new mutations for their evolution. The SP populations began with both the common and rare alleles present at a moment in time in one of the LTEE populations. The MC populations began with six clones with approximately equal initial frequencies. The MP populations started with the most diversity, harboring essentially all of the genetic variation present in the other three treatments at the beginning of the evolution experiment. All 72 populations evolved for 2,000 generations (300 days) in the novel environment, with D-serine as their source of carbon and energy. Using stocks that we froze during the evolution experiment, we subsequently performed competition assays to measure the fitness of the evolved bacteria relative to common competitors, which allowed us to compare the extent of fitness gains among the four treatments. We also tracked a genetic marker embedded in our experiment, which allowed us to observe important dynamics especially during the first 100 generations or so of our experiment.

**FIGURE 1.**
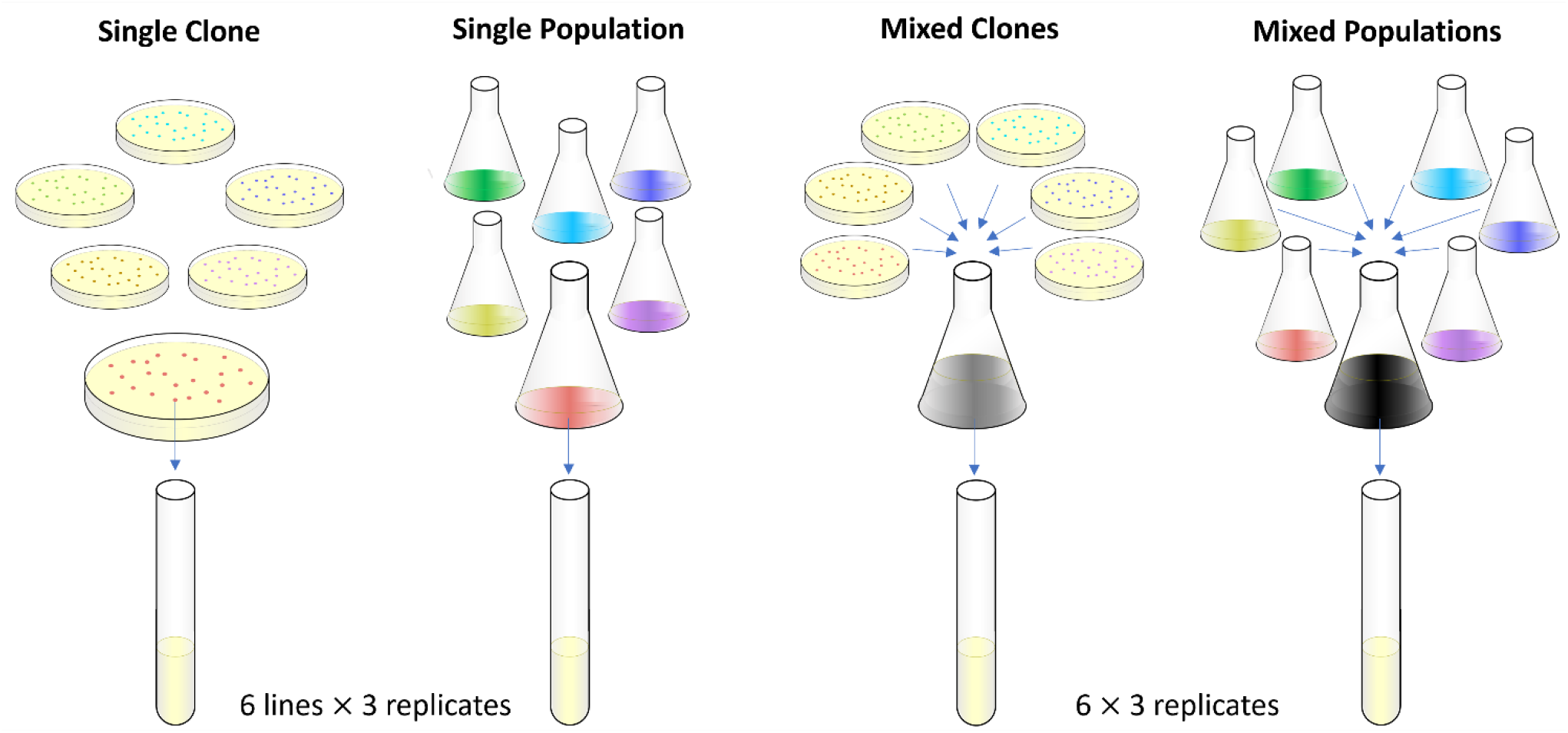
Experimental design. The colors indicate six different founder lineages. The actual colors of colonies on TA indicator agar plates are the same, except the cells derived from the four Ara^−^ lineages produce red colonies while those derived from the two Ara^+^ lineages make pinkish-white colonies. See Materials and Methods for details of the procedures used.

## MATERIAL AND METHODS

### Evolution Experiment in the D-Serine Environment

We used six whole-population samples and six clones from generation 50,000 of the LTEE as the founders for our new evolution experiment (Supplementary File 1). The populations are those named Ara–1, Ara–4, Ara–5, Ara–6, Ara+2, and Ara+5, and from those same populations we used the designated “A” clones that were previously isolated. The whole-population samples and clones were stored at −80°C, where they have remained viable and available for future studies. Two of the six populations (Ara–1 and Ara–4) evolved hypermutability, while the other four retained the low ancestral mutation rate. Before starting our evolution experiment, we re-isolated clones from the freezer stocks for the six A clones on Davis minimal (DM) agar plates supplemented with 4 mg/mL glucose to ensure the genetic homogeneity of the clonal ancestors. Both the re-isolated clones and 120 μL of each whole-population sample were inoculated into 50-mL Erlenmeyer flasks containing 9.9 mL of DM medium supplemented with 1000 μg/mL glucose. These cultures were incubated for 24 h in a shaking incubator at 37°C and 120 rpm. They were then frozen at −80°C with glycerol as a cryoprotectant, in order to generate and preserve samples of the precise ancestral stocks we used for our evolution experiment.

The new evolution experiment itself was begun as follows. On day –2, we inoculated 0.1 mL of each ancestral stock into 9.9 mL of DM medium supplemented with 2000 μg/mL glucose. We incubated these cultures for 24 h in the same conditions as described above. On day –1, we diluted a portion of each culture 100-fold into saline solution, and then transferred 0.1 mL of the diluted culture into 9.9 mL of DM medium supplemented with 25 μg/mL glucose (the same medium as used in the LTEE), and then incubated the cultures for 24 h. On day 0, we took 2 mL from each of the 6 clonal cultures, mixed them well in a flask, and made a starter mix for the MC treatment. We made a similar mix for the MP treatment. We then transferred 0.1 mL of each culture into 9.9 mL of DM medium with 150 μg/mL D-serine (DS150) in an 18 x 150 mm test-tube, vortexed the culture, and then incubated the cultures for 24 h in a standing incubator at 37°C. We prepared 3 biological replicates from each of the 6 clonal and population cultures, making a total of 18 evolving populations in the SC and SP treatments (Figure 1). In those treatments, six sets of three populations shared the same initial genetic background (SC treatment) or the same initial genetic diversity (SP treatment). We also started 18 populations from the clonal starter-mix for the MC treatment, and 18 populations from the population starter-mix for the MP treatment (Figure 1). The 18 populations in the MC treatment share the same set of initial genetic backgrounds, and the populations in the MP treatment share their initial genetic diversity, although very rare alleles might have been distributed unevenly, by chance, among the replicates of these treatments at the start of the evolution experiment.

We transferred the 72 populations (18 populations × 4 treatments) in 9.9 mL of fresh DS150 medium in test-tubes daily, following the same 100-fold dilution protocol for 300 days. In this environment, the populations reach a stationary-phase density of ~5 × 10^7^ cells/mL and total size of ~5 × 10^8^ cells. The bottleneck population size after the 100-fold dilutions is thus ~5 × 10^6^ cells. These values are essentially the same as those for the glucose-limited LTEE populations. We froze samples of each population at −80°C with glycerol as a cryoprotectant every 15 days through day 165, and then every 15 or 30 days through day 300. We also froze the remaining volume of each culture from day 0.

During the evolution experiment, we diluted and spread cells from each population on tetrazolium arabinose (TA) indicator agar plates every 15 days to check for possible cross-contamination among the populations in the SC and SP treatments, where each population derived from either an Ara^−^ or Ara^+^ lineage. We did not find any evidence of contamination during the 300 days of our evolution experiment. The populations in the MC and MP treatments had lineages with both marker states at the start, and we tracked the marker ratio in those populations for evidence of changing ratios, which would indicate fitness differences among the heterogenous founders and their descendants in these populations. To that end, we plated samples from the populations in the MC and MP treatments every other day until day 15, then every three days until day 45, and finally every five days until day 300. There was one interruption in the experiment at day 75. When we restarted the populations from the frozen samples, we plated all of them for the first three days to check whether freezing and reviving the samples altered the relative abundance of the marker states in the MC and MP populations with mixed ancestry. We did not see any substantial changes in the marker ratios, indicating that these steps did not substantially perturb the evolution experiment. Moreover, these procedures were applied to the populations in all four treatments, and thus they would not systematically bias the outcome.

### Fitness Measurements

We isolated clones from each population at generations 500 and 2000 (i.e., days 75 and 300, Supplementary File 1) on DM agar plates with 900 μg/mL D-serine, and we re-streaked the clones on TA plates to confirm their Ara marker state. The clones were chosen at random, except that each clone had the numerically dominant marker state for its source population at these time points for the MC and MP treatments. We then isolated Ara^+^ mutants of several Ara^−^ clones from generation 500 to identify potential common competitors with intermediate fitness relative to other clones from generations 0 to 2000. Using a single pair of common competitors (isogenic except for the Ara marker state) for the fitness assays simplifies procedures and inferences, and having intermediate fitness allows accurate estimates across a wide range of fitness values. We chose MI2228 and an Ara^+^ revertant MI2339 as the common competitors for the main set of fitness assays (Supplementary File 1). MI2228 and MI2339 have equal fitness in DS150 medium, which indicates that the Ara^+^ mutation is selectively neutral in that environment.

On day −2 of the assays, we transferred 0.1 mL from each competitor’s freezer stock into 9.9 mL of Lysogeny Broth (LB) in a 50-mL Erlenmeyer flask, and we incubated the cultures overnight at 37°C and 120 rpm. At day −1, we diluted each culture 100-fold in saline solution, transferred 0.1 mL into 9.9 mL of DS150 medium in a test-tube, vortexed it, and then incubated the cultures for 24 h in a standing incubator at 37°C. This day served as the conditioning step to ensure that competitors were acclimated to the environment where they would compete, and where the experimental populations had evolved. The rest of the procedure is the same as described elsewhere for the LTEE (Lenski et al., 1991; Wiser et al., 2013), except for the medium and culture vessel. In brief, we always competed the common competitor with the opposite marker state from the clone of interest. We transferred 0.05 mL of each competitor’s acclimated culture into 9.9 mL of DS150, and vortexed the new culture to mix the two competitors. We immediately took a sample, diluted it in saline solution, and plated cells on TA agar. The cultures were incubated for 24 h, at which time we again sampled the cultures and plated cells on TA agar. The resulting red (Ara^−^) and white (Ara^+^) colonies were counted after the plates were incubated for a day at 37°C. We calculated each competitor’s realized growth rate as the log-transformed ratio of its final and initial densities. We then computed the fitness of the clone of interest relative to the common competitor as the ratio of their growth rates during the competition.

We used the generation 0 stocks multiple times for estimating initial fitness levels. We have only 12 generation 0 stocks because we used the same six clones for the three replicates of each clone in the SC treatment, and the same six whole-population samples for the three replicates of each population in the SP treatment. The populations in the MC and MP treatments were derived from their respective starter mixes. We cannot measure the fitness of samples that contain both Ara^−^ and Ara^+^ cells using our method, which relies on a common competitor with the opposite marker state. Therefore, we used the same six clonal stocks at generation 0 for both the SC and MC treatments, and the same six population stocks at generation 0 for both the SP and MP treatments, and for all three replicates.

We also ran a second set of competition assays using the LTEE ancestors, REL606 and REL607, as common competitors. For these assays only, we used a 1:4 starting ratio at day 0, instead of the 1:1 starting ratio described above, because of the substantially lower fitness of the LTEE ancestors in comparison to the common competitors used above. Specifically, we began each competition assay by mixing 0.08 mL of REL606 or REL607 and 0.02 mL of the strain of interest in the test-tube containing the DS150 medium. The assay conditions and the calculations of relative fitness were otherwise the same.

### Statistical Analyses

All of our statistical analyses were performed using the referenced tests in R version 4.2.0. The analysis scripts and underlying data will be deposited in the Dryad Repository upon acceptance of this paper.

## RESULTS

### Effect of Initial Variation on Fitness Improvement

To assess the effect of the initial within-population diversity on adaptation to the new environment, we measured the relative fitness of the evolved bacteria by competing them against the common competitor strains. We cannot measure fitness of the entire evolved populations using our method, however, because that method requires mixing the evolved bacteria with the common competitor strain bearing the alternative Ara marker, and some populations in the MC and MP treatments had descendants of lineages with both marker states. Therefore, we isolated random clones at generations 500 and 2000 as representatives of each population, and we measured their fitness. For generation 0, we used the stocks of the founder clones and populations that we froze immediately after the start of the evolution experiment. We used the six clone stocks that we had used to found populations in the SC and MC treatments as the generation 0 samples for those treatments, and we used the six whole-population stocks used to found populations in the SP and MP treatments as the generation 0 samples for those treatments. As a consequence, the generation 0 samples for the SC and MC treatments are technically identical, as are those for the SP and MP treatment.

Figure 2 shows the trajectories of the ln-transformed relative fitness values for the four treatments. As a reminder, the replicate populations in the SC and SP treatments had six different founding backgrounds. In contrast, the replicate populations in the MC and MP treatments originated from the same starter mix of six clones or six whole populations, respectively, and thus the replicates in those treatments shared the same founding backgrounds and diversity. The rate of increase in relative fitness clearly slowed over time in the D-serine environment (Figure 2). That deceleration is similar to what was seen during the first 2,000 generations in the glucose-limited environment of the LTEE (Lenski et al., 1991), and it is indicative of diminishing-returns epistasis (Wiser et al., 2013).

**FIGURE 2.**
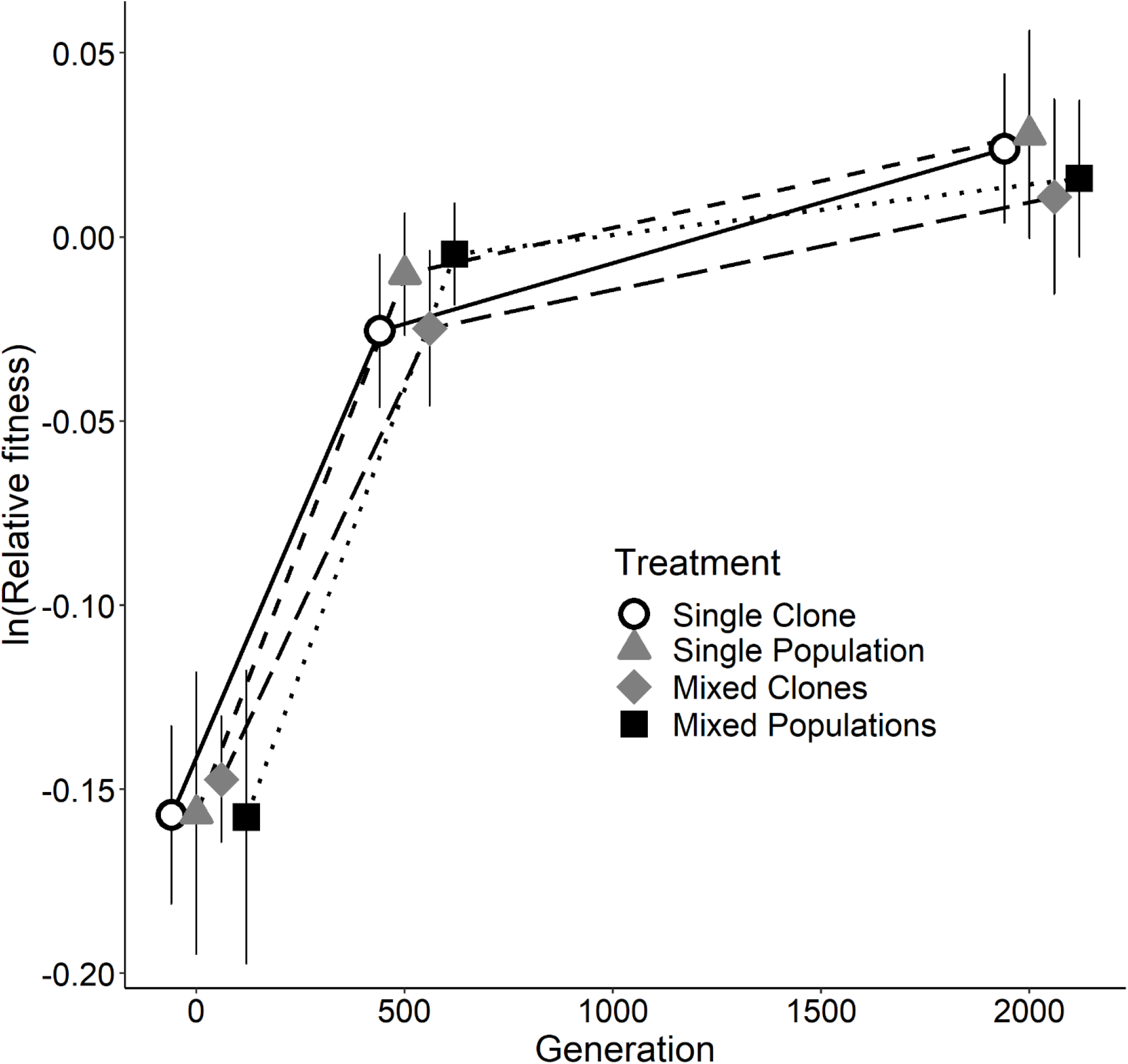
Relative fitness of the four treatments at generations 0, 500, and 2000 in the D-serine environment. Each symbol is the mean of 18 ln-transformed fitness estimates, and error bars show 95% confidence intervals. See Figure S1 for each replicate population.

Most importantly for our aims and questions, we found no significant difference in fitness among the four treatments at either generation 500 or 2000 (*p* = 0.2300 and *p* = 0.7213, respectively; one-way ANOVA, Table S1). The absence of meaningful differences among the treatments in the rate and extent of their adaptation was surprising to us, given the different levels of within-population genetic diversity at the beginning of the experiment. One possible explanation for the negative results with respect to differences in the final fitness values is that the initial variation present in treatments SP, MC, and MP did not include alleles that were sufficiently beneficial in the novel environment relative to new mutations. In other words, the populations in all four treatments ultimately depended on new mutations for adaptation to the novel D-serine medium, regardless of the different levels of initial genetic diversity. In the sections that follow, we present and examine additional data that helps to explain this result.

### Marker Trajectories During the Evolution Experiment

We tracked the relative abundance of the two Ara marker states in all treatments during the evolution experiment (Figures 3 and S2). The populations in the SC and SP treatments began with a single marker state; in these populations, checking the marker states allowed us to check for cross-contamination, which we did not see. The populations in the MC and MP treatments began with a mix of the two marker states. By tracking the relative abundance of the two states in those populations, we could observe the effects of both initial fitness variation linked to the markers and later beneficial mutations that gave rise to selective sweeps. The MC and MP treatments started with equal culture volumes of four Ara^−^ lineages and two Ara^+^ lineages; therefore, the log-transformed ratios of Ara^−^ to Ara^+^ cells were initially > 0 for all of the populations in those treatments (Figures 3 and S2).

**FIGURE 3.**
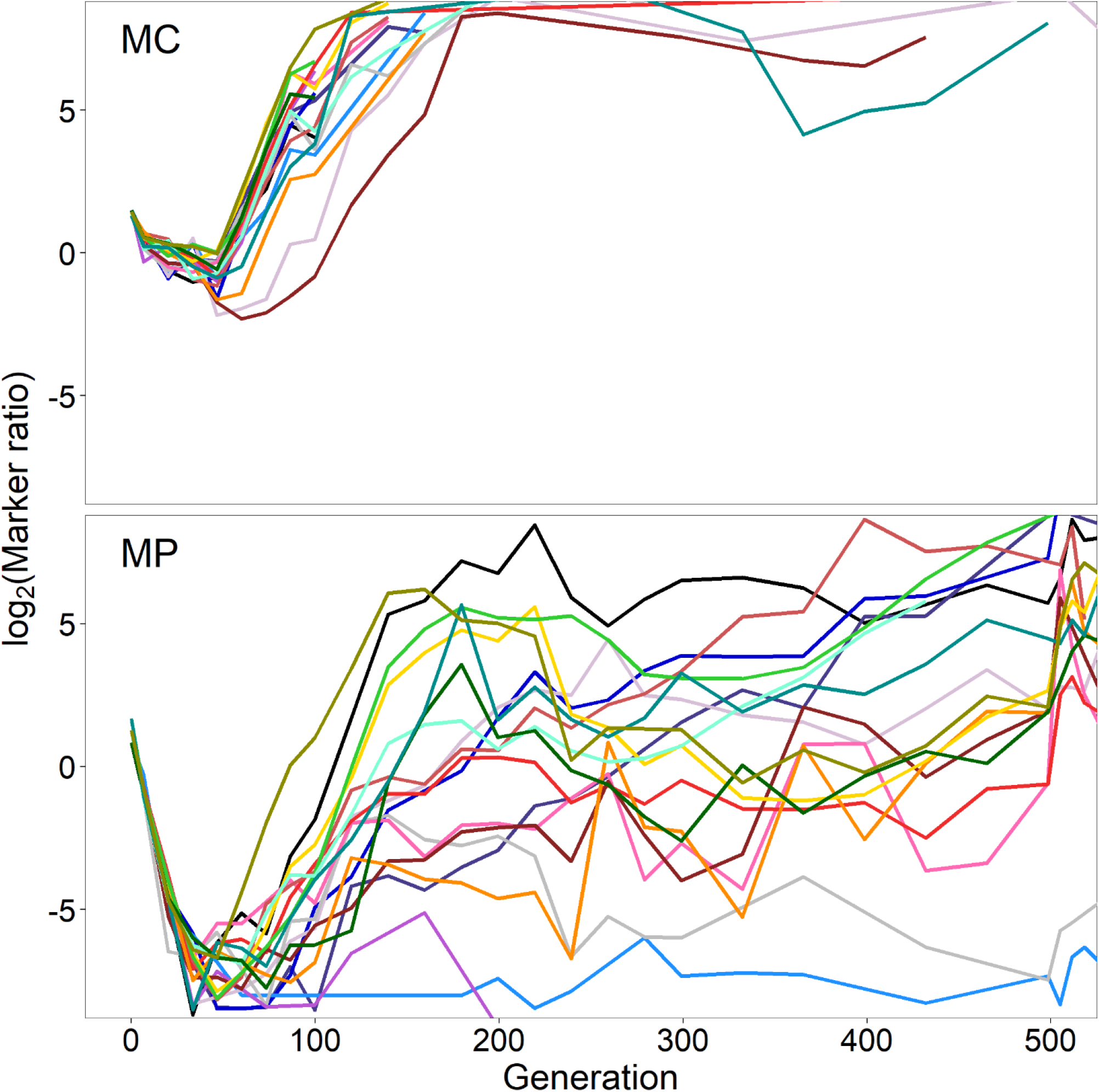
Marker trajectories in the Mixed-Clones (**MC**) and Mixed-Populations (**MP**) treatments during the first 500 generations. The marker ratio indicates the number of cells derived from the Ara^−^ founder lineages divided by the number of cells derived from the Ara^+^ founder lineages.

We observed strikingly similar marker trajectories among the 18 replicate populations in the MC and MP treatments, especially during the first ~100 generations (Figures 3 and S2). Despite the initially greater number of Ara^−^ lineages, cells derived from one or more Ara^+^ lineages increased in relative abundance in all 36 populations. By 30-50 generations, the Ara^+^ cells were numerically dominant in all 18 MP populations and in most MC populations as well (Figure 3). These initial “bursts” imply that one or more of the Ara^+^ clones and populations initially present in the MC and MP treatments were substantially more fit than the Ara^−^ clones and populations. We will return to this point in the next section.

By generation 100, all 18 populations in the MC treatment, and most of the MP populations, had reversed course, with descendants of one or more Ara^−^ lineages rising sharply in abundance relative to the Ara^+^ descendants (Figure 3). The Ara^−^ descendants remained numerically dominant through the first 500 generations in all 18 MC populations (Figure 3, top), and they evidently fixed in all 18 cases by 2,000 generations (Figure S2, top). By contrast, the later marker-ratio trajectories of the MP populations were much more variable. Descendants of Ara^−^ founders were usually more abundant through the first 500 generations, but with tremendous dispersion between the trajectories (Figure 3, bottom). By 2,000 generations, most MP populations had also evidently fixed one of the marker states, but with several fixations in each direction (Figure S2, bottom).

The marker-ratio trajectories also show that bursts leading to the early rise of cells derived from one or more Ara^+^ lineages were much steeper for the populations in the MP treatment than for those in the MC treatment. While the initial ratios were virtually identical, at generation 47 (day 7) the mean log2 ratios were –0.825 and –7.004 for the MC and MP treatments, respectively, even excluding two MP populations without any Ara^−^ cells among the hundreds sampled. In fact, all 18 MP populations had a much lower ratio than any of the 18 MC populations, a difference that is highly significant (*p* << 0.0001; two-tailed Welch’s *t*-test). We chose day 7 for this comparison because that is when the MC treatment showed the lowest average log ratio, although several adjacent days show a similarly stark difference between these two treatments.

In summary, we observed strikingly similar marker trajectories among the replicate populations in the MC and MP treatments in the early generations of our evolution experiment. Given the inevitable genetic linkage in asexual populations, this pattern implies that the metagenomes of the populations also evolved in parallel during this early phase. Moreover, this parallelism indicates that selection acted on shared genetic variation present in these populations at the start of experiment (i.e., identical by descent). It is reminiscent of the repeatability observed in previous evolution experiments with other organisms that were also founded by populations with shared initial variation (Burke et al., 2014).

### Fitness Differences Among the Founder Clones and Populations

We examined the relative fitness values of the six founding populations and the six founding clones to better understand the similar early marker trajectories seen among the replicate populations in the MP and MC treatments, as well the difference between those treatments in the slope of those early trajectories (Figure 3). For these analyses, we use the same data as the generation 0 data that underlies the grand means for each treatment in the fitness trajectories (Figure 2).

Given the consistent marker-ratio trajectories towards the Ara^+^ marker state, we expect to see that one or both of the Ara^+^ founders had the highest fitness. Also, given that the early trend toward the Ara^+^ state was much faster in the MP treatment than in the MC treatment, we expect that fitness differential to be greater among the whole-population founders than among the clonal founders. Figure 4 shows the relative fitness of the founding populations (panel A) and clones (panel B). In each panel, note that we have arranged the founders from the lowest to highest relative fitness.

**FIGURE 4.**
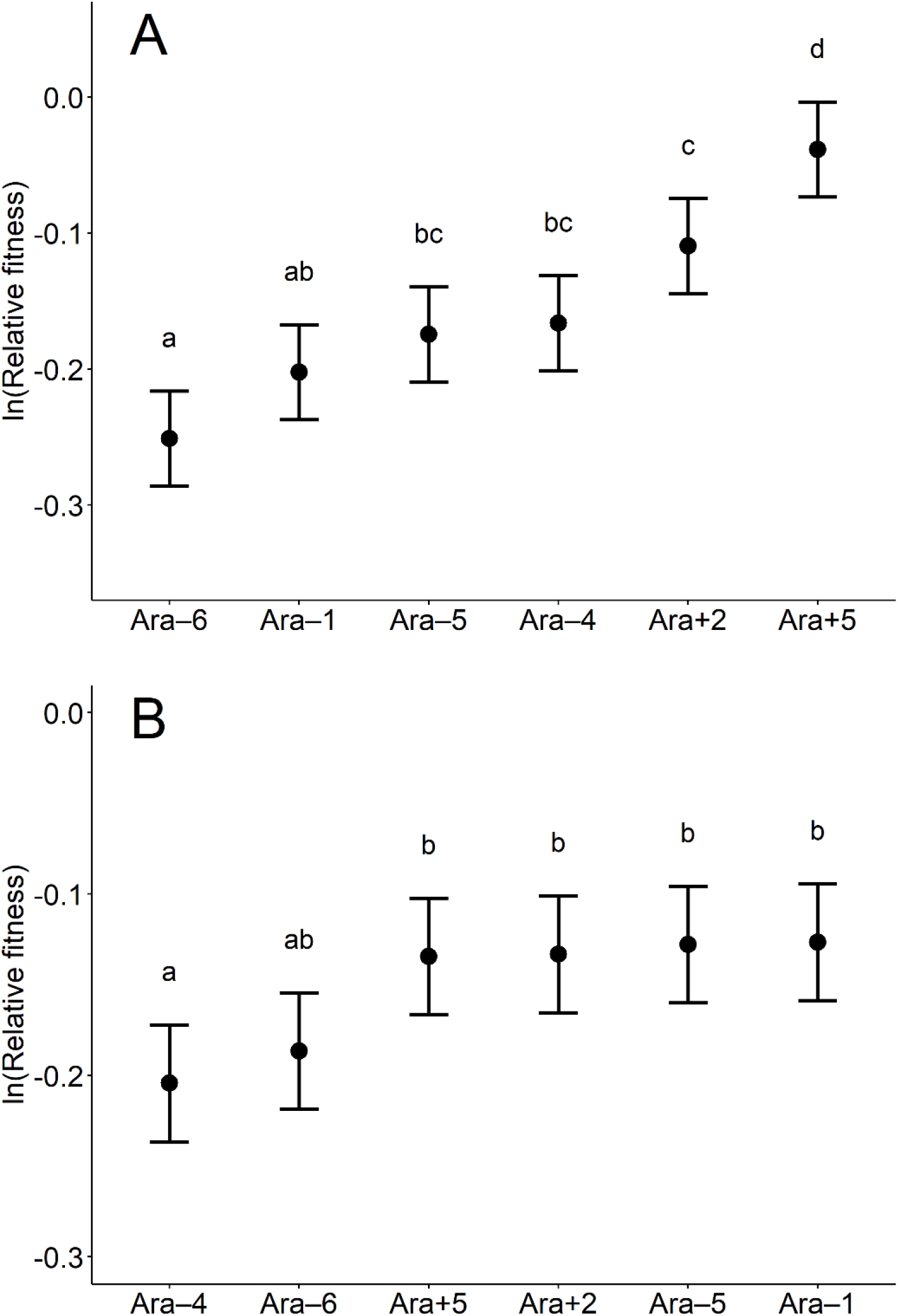
Relative fitness of founder whole populations (**A**) and founder clones (**B**). The founders in each panel are arranged from lowest to highest fitness. The filled circles show the mean value of the ln-transformed fitness, based on 18 replicates for each founder. The error bars show 95% confidence limits, based on the *t*-distribution with 17 degrees of freedom and using the pooled standard deviation estimated from the corresponding ANOVAs (Table S2). Letters above the error bars identify sets of founders with values that are not significantly different, based on Tukey’s test for multiple comparisons (*p* > 0.05). For this analysis, we combined data for the SC and MC treatments, and similarly we combined data for the SP and MP treatments, because we used the same 6 clonal or whole-population samples at generation 0 for those pairs of treatments (see Materials and Methods).

Focusing first on the whole-population data (Figure 4A), we see that both of the Ara^+^ founders have higher mean fitness than any of the Ara^−^ founders in the DS150 environment. An ANOVA confirms that there is significant variation in fitness among the founders (*p* < 0.0001, Table S2, top), and Tukey’s test confirms that the Ara+5 whole-population founders are significantly more fit than any of the Ara^−^ founders. These results thus support our expectation from the marker trajectories that one or both of the Ara^+^ founders had the highest fitness.

When we look at the corresponding data for the clonal founders, we see a more ambiguous pattern (Figure 4B). The relative fitness levels of the clones are more similar; four clones (two Ara^+^ and Ara^−^) are virtually identical to one another and slightly higher than two others (both Ara^−^). An ANOVA confirms that there is significant difference in fitness among the clone founders (*p* = 0.0004, Table S2, bottom), while Tukey’s test finds no significant difference in fitness among the several most fit founder clones.

Based on the ANOVAs, we estimated the among-founder variance components, *V_A_*, for fitness in these two treatments (Sokal and Rohlf, 1995). That founding variation is what would fuel the earliest response to selection in the evolution experiment before new mutations have had enough time to become relevant. As expected, the estimated variance in fitness among the whole-population founders (*V_A_* = 0.0052) is much greater than among the clonal founders (*V_A_* = 0.0009).

We also performed an additional set of competition assays to estimate the fitness of the founders of our evolution experiment relative to a different pair of common competitors. In this case, we competed the six founders of whole-populations and clones against the marked ancestors of the LTEE (Figure S3). The founders generally had higher fitness relative to the LTEE ancestors than relative to the common competitors used in our other assays. Therefore, we used a 1:4 starting ratio of the founders relative to the LTEE ancestors, instead of the 1:1 starting ratio used in the other competitions (Materials and Methods). Otherwise, the assay conditions and calculations of relative fitness as the ratio of realized growth rates were the same. We also arranged and analyzed these data as before.

These additional data also support one of our two expectations based on the marker trajectories, namely, that one Ara^+^ founder had higher fitness than any of the Ara^−^ founders. In this case, we see that Ara+5 has the highest mean fitness among both the whole-population (Figure S3A) and clonal (Figure S3B) founders. The results of the Tukey tests confirm that Ara+5 had significantly higher fitness than all other whole-population founders and higher fitness than all but one clonal founder. The ANOVAs indicate significant variation in fitness among both the whole-population (Table S3, top) and clonal (Table S3, bottom) founders. However, the variation in fitness is not greater among the whole-population founders than among the clonal founders. The estimated among-founder variance component for fitness for the whole-population founders (*V_A_* = 0.0274) is essentially identical to the variance among the clonal founders (*V_A_* = 0.0269).

Across the four sets of competitions (founder clones and whole populations, against two pairs of common competitors), we find that the founders derived from LTEE population Ara+5 had the highest fitness in three of these sets (Figures 4A, S3A, and S3B), while they were tied for the highest fitness in one set (Figure 4B). These results clearly imply that the early trends toward the Ara^+^ state in the marker-ratio trajectories in the MC (Figure 3, top) and especially the MP (Figure 3, bottom) treatments were caused by the initial fitness advantage that the Ara+5 founders had in the new DS150 environment. By contrast, the subsequent reversals in most trajectories are presumably associated with new mutations that arose during our evolution experiment. (In theory, very gradual and uniform reversals could occur even without new mutations if the single most fit founder had a different marker state than the maker state with the higher average fitness across its constituent lineages. However, this hypothetical scenario is clearly not the case for the MP treatment, nor can it explain the variation in the time and strength of the reversals in the MP and MC treatments shown in Figure 3.) A deeper understanding of the reversals will require future genomic analyses, as we explain in the Discussion.

## DISCUSSION

It is generally difficult to disentangle the role of standing genetic variation and new mutations in the process of adaptation by natural selection. Even with experiments, different study systems tend to emphasize one source or the other. Selection experiments that use sexually reproducing plants and animals have typically started from base populations that harbor substantial standing variation, and they rarely run for more than a few tens of generations owing to the long generation time of these organisms. As a consequence, these experiments rely largely on variation that was present at the start of the experiment to fuel the response to selection. The field of experimental evolution with bacteria and other microorganisms has expanded greatly in recent years (Barrick and Lenski, 2013; Lenski, 2017b; Van den Bergh et al., 2018). These study organisms have rapid generations, and most of them reproduce asexually during the experiments, even those that may undergo parasexual recombination (e.g., horizontal gene transfer) in nature. Our experiment was designed to compare the contributions of initial genetic variation and new mutations during adaptation of strictly asexual populations to a new environment.

To that end, we constructed four treatments with different initial levels of genetic diversity. Each treatment had 18 populations. In all cases, the founders came from the LTEE, in which *E. coli* have evolved in and adapted to a glucose-limited medium for 50,000 generations. At one extreme, each new population was founded by a single genotype, and thus there was no initial within-population diversity. We call this the Single-Clone (SC) treatment; six different clones, each derived from a different LTEE lineage, were used to found three replicate populations. At the other extreme, 18 populations derived from an admixture of six whole-population samples that included both common and rare genotypes from the source populations. We call this the Mixed-Populations (MP) treatment. We also had two treatments that started with intermediate levels of genetic variation, which we call the Single-Population (SP) and Mixed-Clones (MC) treatments (Figure 1).

We propagated all 72 populations for 2000 generations in a new environment, one in which D-serine replaced glucose as the source of carbon and energy. We then measured the fitness of evolved strains from each population at both 500 and 2000 generations. We observed rapid early adaptation to the D-serine environment in all of the populations, but the rate of further fitness improvement declined over time, similar to what has been seen in the glucose environment of the LTEE (Wiser et al., 2013) as well as seen in other microbial evolution experiments (e.g., Johnson et al., 2021; Marad et al., 2018).

Most importantly, however, we found no significant differences among the four treatments in their mean fitness at generations 500 and 2000 (Figure 2), despite their different levels of genetic diversity at the beginning of the experiment. Thus, the populations in the SC treatment, each of which had no genetic diversity at the start, achieved the same fitness as the populations in the MP treatment, which started with all the diversity found in six LTEE populations combined. One possible explanation for this negative result would be that there were simply no differences in fitness in the D-serine medium among the founders in the treatments that began the experiment with genetic variation. In that case, all the populations in all four treatments would have had to depend entirely on new mutations to fuel adaptation to the new medium. But as we discovered, there was significant initial within-population variation for fitness in the new environment, at least in the MC and MP treatments.

Our first evidence of that initial fitness variation came from tracking the ratio of a neutral genetic marker that differed among the LTEE-derived founders, and which was therefore polymorphic in each of the populations in the MC and MP treatments. If there was no initial fitness variation in the new environment, then that ratio should have remained constant (within sampling error) until such time as a beneficial mutation occurred and began to sweep through one or the other marked backgrounds, thereby perturbing that ratio (Barrick et al., 2010; Izutsu et al., 2021). Alternatively, if the different founding genotypes had unequal fitness, then the marker ratio would systematically and immediately deviate from its initial value as a result of the inevitable linkage in asexual genomes between the marker and the alleles responsible for the fitness differences. This alternative outcome is precisely what we saw. We observed strikingly similar directional shifts in the marker-ratio trajectories among populations in the MC and MP treatments, especially during the first ~100 generations (Figure 3). These parallel directional trajectories imply the presence of at least one “preadapted” genotype among the founders in those treatments.

We also compared the relative fitness of the founding clones and founding populations used in the MC and MP treatments, respectively. These comparisons showed that the founders derived from LTEE Ara+5 lineage had fitness as high as or higher than the other founders in the new D-serine environment (Figure 4 and S3), consistent with the early and systematic shifts in the marker-ratio trajectories to the Ara^+^ marker state. Also, the early marker-ratio trajectories in the MP treatment were much steeper than in the MC treatment (Figure 3), consistent with greater fitness differentials favoring the Ara+5 founders in the MP treatment (Figure 4). Thus, the genetic variation initially present in the MC and MP populations drove adaptation to the new environment during the first 100 generations of our experiment. However, new beneficial mutations soon arose that perturbed and often reversed those early trends in the marker ratios (Figure 3). By generation 500, the beneficial effects of these new mutations were sufficiently large that the initial variation no longer mattered, and all four treatments—including even the SC treatment, in which each population started from a single clone—had achieved similar average fitness (Figure 2).

One might have expected that new beneficial mutations would have arisen randomly with respect to the marker state of the founders in the MC and MP treatments. Four of the six founders came from LTEE lineages with the Ara^−^ marker state, and two from lineages with the Ara^+^ marker state. If the mutations that were beneficial in the D-serine environment arose very early in the new experiment, then we might expect about two-thirds of the marker trajectories to reverse course and trend toward the Ara^−^ state, after those mutations reached high frequency within the Ara^−^ subpopulation. The expected fraction might be lower than two-thirds, however, because the Ara^+^ subpopulation was increasing in frequency, and would be expected to generate an increasing proportion of the beneficial mutations, all else being equal. Contrary to this naïve expectation, however, all 18 populations in the MC treatment and 15 of the 18 populations in the MP treatment ended the experiment with descendants of the Ara^−^ founders being numerically dominant (Figure S2).

This bias implies that one or more of the Ara^−^ founders had greater potential for future adaptation than other founders. Of the six LTEE lineages that provided the founders used in our study, two of them—both with the Ara^−^ state—evolved hypermutability during the LTEE (Tenaillon et al., 2016). The Ara–4 lineage became defective in mismatch repair (Sniegowski et al., 1997), while the Ara–1 lineage acquired mutations in two enzymes that would normally prevent the misincorporation of oxidized nucleotides into DNA (Wielgoss et al., 2013). It is also possible that epistasis between new mutations and the various genetic backgrounds has led to differences in evolvability among the various founders. Background-dependent epistasis leading to differences in evolvability has been observed in the LTEE using replay experiments (Woods et al., 2011; Blount et al., 2012; Wünsche et al., 2017). In any case, the populations in the MC and MP treatments had reached similar fitness levels to those in the SC and SP treatments by generations 500 and 2000. Thus, the effects of both the initial standing variation and differences among the founders in their genetic potential for adaptation impacted only the earliest phases of evolution in the new D-serine environment.

Genetic variation is essential for populations to adapt to a new environment. We observed that pre-existing variation was important during the first ~50 generations in the D-serine medium, leading to substantial changes in the relative abundance of the different founders in the MC and MP treatments (Figure 3). Those changes depended on the initial genetic variation, which was identical by descent across the replicate populations in those treatments, and thus they indicate collateral evolution (Stern, 2013; Lenski, 2017a). By contrast, the subsequent reversals in the relative abundance of descendants of those founders, and the fact that populations in all four treatments eventually achieved similar fitness levels (Figure 2), resulted from new mutations that arose independently in those populations, indicating parallel evolution (Stern, 2013; Lenski, 2017a). Thus, we observed both collateral and parallel evolution in our experiment with bacteria.

Two long-term experiments using *Drosophila* also reported collateral evolution, but they were not followed by parallel evolution (Burke et al., 2010; Graves et al., 2017). The longer generations and smaller populations of fruit flies probably limited the supply of new beneficial mutations, while sexual reproduction and the resulting segregation of pre-existing variation may have continued to fuel the ongoing response to selection. The importance of sexual reproduction with respect to the contributions of collateral versus parallel evolution was also evident in an evolution experiment performed using yeast (Burke et al., 2014). That experiment ran for 540 generations with large populations (10^6^ cells during the transfer bottlenecks), and the populations were founded by a diverse set of diploids obtained by crossing wild strains. Although yeast can reproduce asexually, they underwent periodic mating and recombination in their experiment. As a consequence, segregating variation derived from the founders evidently fueled adaptation for the duration of the experiment, with little or no input from new beneficial mutations (Burke et al., 2014).

In any case, our study has shown that strictly asexual populations can also benefit from pre-existing variation, but the effect is likely to be smaller than in sexual populations. Moreover, any benefit of pre-existing variation in asexual populations may often be short-lived, as we saw in our experiment, because that variation will be purged when new beneficial mutations sweep to fixation. In particular, it appears that the pre-existing alleles provided by the founders in our study were not sufficiently beneficial in the D-serine environment, such that the populations readily produced new mutations that provided greater benefits and displaced the initial variants. Even the populations in the SC treatment, which had no genetic diversity at the start of the experiment, achieved fitness levels comparable to the other treatments (Figure 2).

In future work, we intend to sequence the genomes of the founders and evolved samples from several timepoints. In addition to shedding light on the genetic basis of adaptation to growth on D-serine, genomic data will enable us to test and refine our inferences based on the fitness measurements and marker-ratio trajectories. In particular, we make several predictions that can be tested using genomic data. First, we expect to find an increased frequency of diagnostic alleles from the Ara+5 founders in the early (~50 generations) metagenomic samples from all of the populations in the MC and MP treatments. Second, we expect to see the alleles from Ara+5 subsequently disappear in all MC and most MP populations. Third, we predict that diagnostic alleles from one or more of the Ara^−^ founders will achieve numerical dominance in all of the MC and many MP populations by generation 500 and remain dominant through generation 2000. In addition, genomic data should clarify whether one or both of the hypermutable founders (Ara–1 and Ara–4) in the MC and MP treatments dominated over time in a manner consistent with their having greater evolvability, in the sense of being able to adapt to the new environment. If so, that raises the interesting question of how the populations derived from the non-mutator founders in the SC and SP treatments achieved similarly high fitness. Perhaps, for example, the populations founded by mutators and non-mutators had similar beneficial mutations, but the hypermutators acquired them slightly earlier in the experiment.

In closing, our study contributes to filling the gap between the different experimental designs that are typically used with different model systems, and to understanding how these differences impact the dynamics and repeatability of evolution. While it remains difficult to observe adaptation driven by new mutations using long-standing model systems like *Drosophila*, we demonstrate that one can disentangle and estimate the contributions of standing variation and new mutations to adaptation in microbial systems. We also show that these contributions may depend on the particular history of the founders, and that the relative contributions of pre-existing variation and new mutations are highly sensitive to when they are measured after the evolving populations encounter a new environment.

## ACKNOWLEDGMENTS

We thank Sydney Anway, Kate Bellgowan, Devin Lake, and Zachary Matson for assistance in the lab, and Thomas LaBar for valuable discussion. We acknowledge funding from the National Science Foundation (DEB-1813069), the USDA National Institute of Food and Agriculture (MICL02253), the BEACON Center for the Study of Evolution in Action (NSF Cooperative Agreement DBI-0939454), the Yamada Science Foundation (to MI), and the Japan Society for the Promotion of Science (to MI).

