## Supplementary Tables and Figures for "Experimental Test of the Contributions of Initial Variation and New Mutations to Adaptive Evolution in a Novel Environment"

### SUPPLEMENTARY TABLES

**TABLE S1 |** One-way ANOVAs of ln fitness at generations 0, 500, and 2000

Generation 0

| Source | DF | SS | MS | *F* | *p* |
|---|---|---|---|---|---|
| Treatment | 3 | 0.0013 | 0.0004 | 0.1087 | 0.9547 |
| Error | 68 | 0.2690 | 0.0040 | | |
| Total | 71 | 0.2703 | | | |

Generation 500

| Source | DF | SS | MS | *F* | *p* |
|---|---|---|---|---|---|
| Treatment | 3 | 0.0059 | 0.0020 | 1.4716 | 0.2300 |
| Error | 68 | 0.0915 | 0.0013 | | |
| Total | 71 | 0.0975 | | | |

Generation 2000

| Source | DF | SS | MS | *F* | *p* |
|---|---|---|---|---|---|
| Treatment | 3 | 0.0032 | 0.0011 | 0.4454 | 0.7213 |
| Error | 68 | 0.1612 | 0.0024 | | |
| Total | 71 | 0.1644 | | | |

**TABLE S2 |** ANOVAs of founders' fitness relative to the common competitors

Whole-population founders

| Source | DF | SS | MS | *F* | *p* |
|---|---|---|---|---|---|
| Strain | 5 | 0.4967 | 0.0993 | 20.0850 | < 0.0001 |
| Error | 102 | 0.5045 | 0.0049 | | |
| Total | 107 | 1.0012 | | | |

Clone founders

| Source | DF | SS | MS | *F* | *p* |
|---|---|---|---|---|---|
| Strain | 5 | 0.1049 | 0.0210 | 5.0032 | 0.0004 |
| Error | 102 | 0.4276 | 0.0042 | | |
| Total | 107 | 0.5325 | | | |

**TABLE S3 |** ANOVAs of founders' fitness relative to another pair of common competitors

Whole-population founders

| Source | DF | SS | MS | *F* | *p* |
|---|---|---|---|---|---|
| Strain | 5 | 0.4547 | 0.0909 | 10.4428 | 0.0005 |
| Error | 12 | 0.1045 | 0.0087 | | |
| Total | 17 | 0.5592 | | | |

Clone founders

| Source | DF | SS | MS | *F* | *p* |
|---|---|---|---|---|---|
| Strain | 5 | 0.4332 | 0.0866 | 14.6447 | < 0.0001 |
| Error | 12 | 0.0710 | 0.0059 | | |
| Total | 17 | 0.5042 | | | |

### SUPPLEMENTARY FIGURES

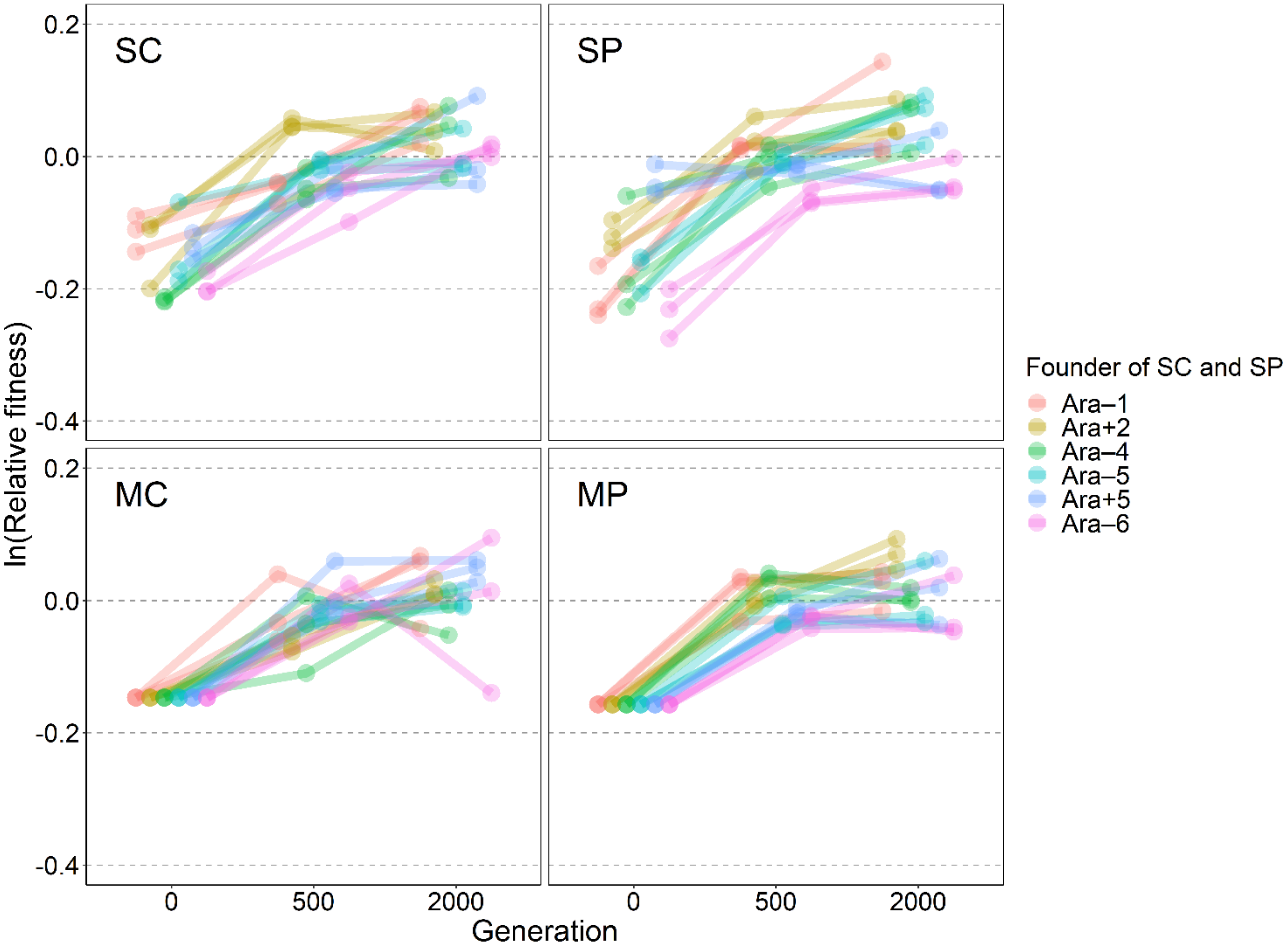


**FIGURE S1 |** Mean fitness for each experimental population in the four treatments. Each point shows the mean of three replicate assays. Figure 2 shows the grand mean of the 18 means shown here for each timepoint and treatment. The colors indicate the founder LTEE strains for the Single-Clone (**SC**) and Single-Population (**SP**) treatments. Although there is no difference in the founders used for the 18 populations in the Mixed-Clones (**MC**) and Mixed-Populations (**MP**) treatments, we use the same color scheme for those treatments in order to distinguish the populations at later generations. Recall that fitness values for generation 0 of the MC treatment, and similarly for the MP treatment, were measured using the same samples as those for the SC and SP treatments, respectively (see Materials and Methods). The 18 populations in those treatments derived from the same starter mixes, and thus all 18 had the same fitness at generation 0, which we calculated as the grand mean.

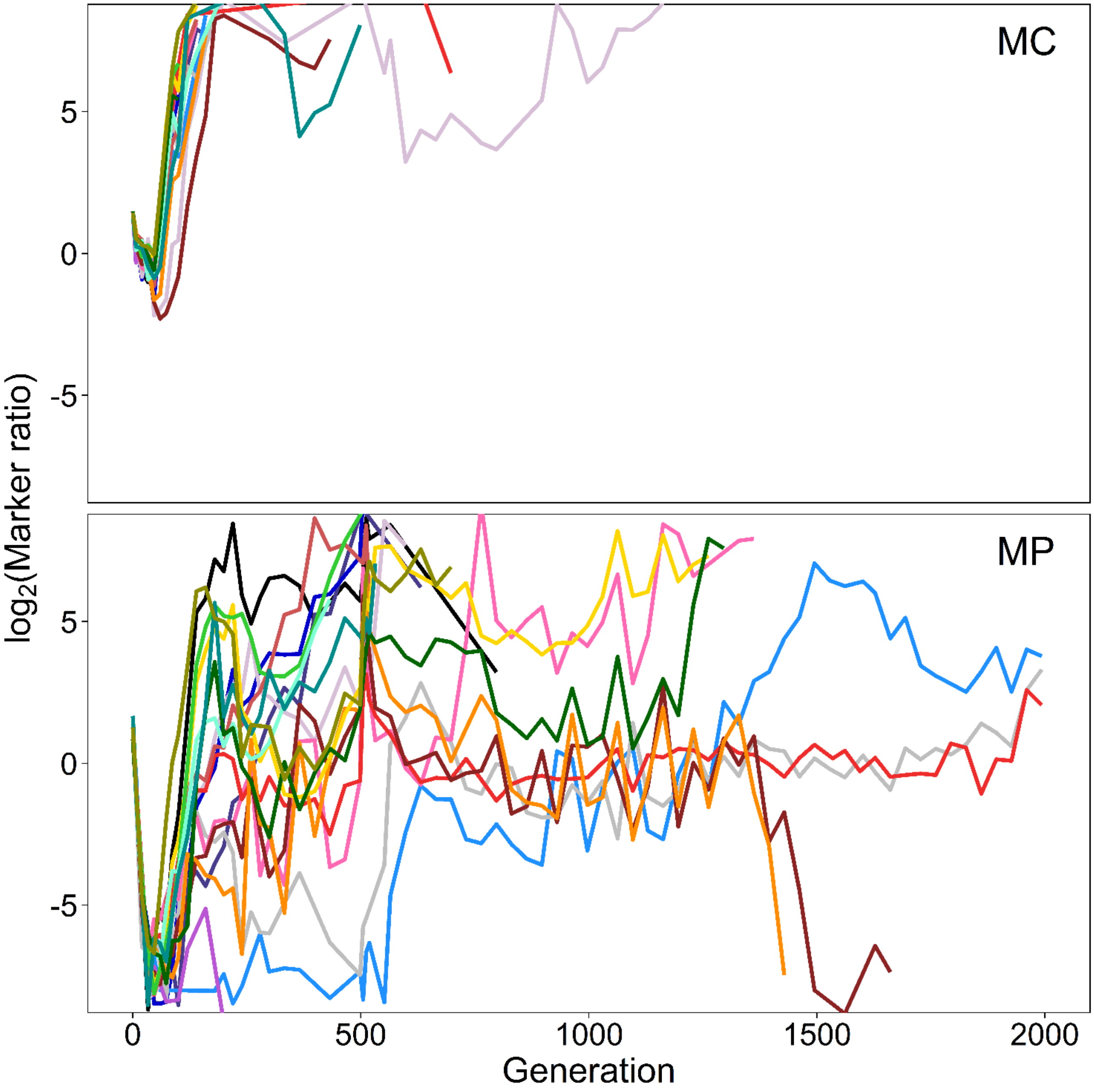


**FIGURE S2 |** Marker trajectories in the Mixed-Clones (**MC**) and Mixed-Populations (**MP**) treatments during the 2000 generations of the evolution experiment. The marker ratio indicates the number of cells derived from the Ara$^{-}$ founder lineages divided by the number of cells derived from the Ara$^{+}$ founder lineages.

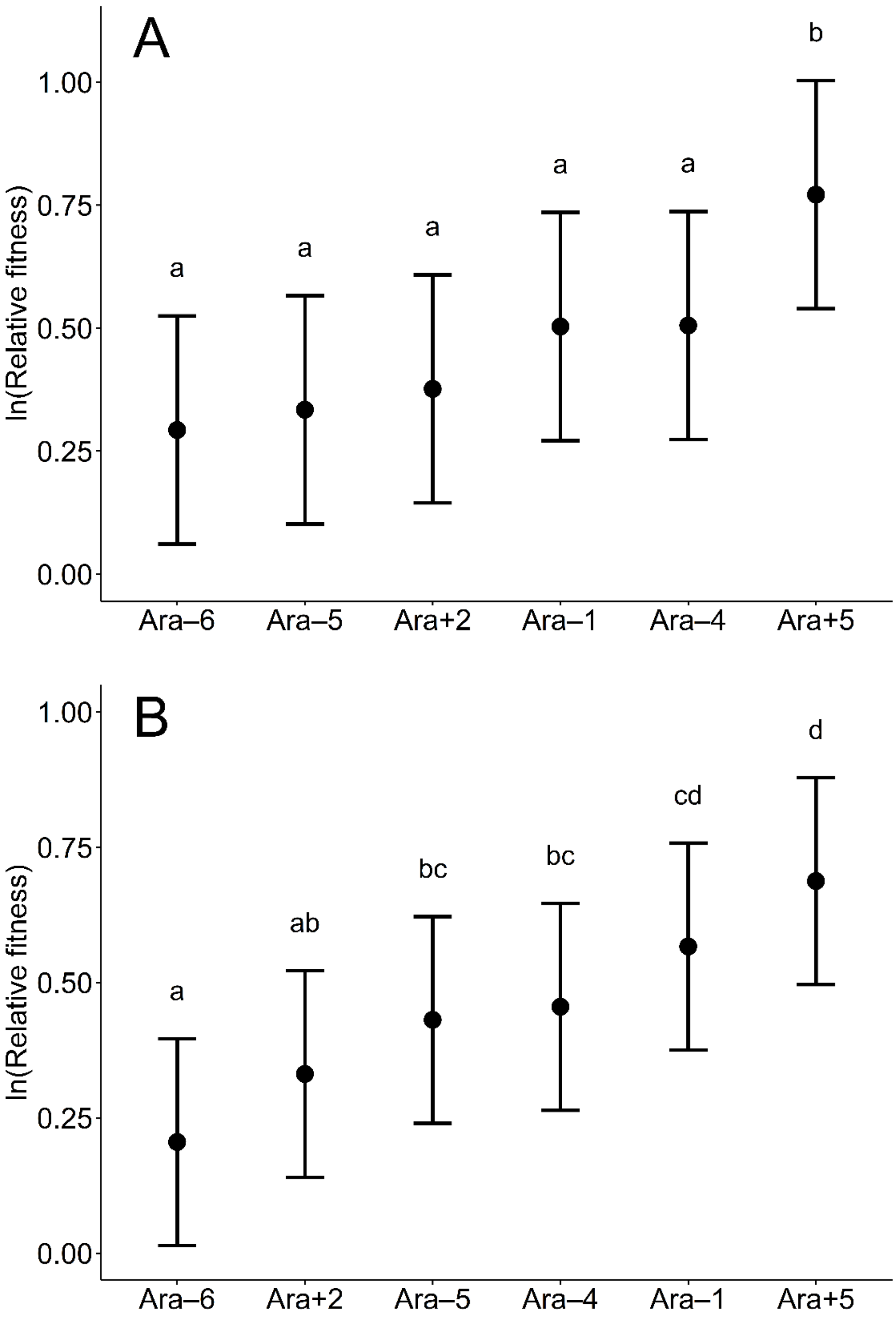


**FIGURE S3 |** Relative fitness values of founder whole populations (**A**) and founder clones (**B**) relative to the ancestors of the LTEE. The founders in each panel are arranged from lowest to highest fitness. The filled circles show the mean value of the ln-transformed fitness, based on 3 replicates for each founder. Error bars show 95% confidence limits, based on the *t*-distribution with 2 degrees of freedom and using the pooled standard deviation estimated from the corresponding ANOVAs (Table S3). Letters above the error bars identify sets of founders with values that are not significantly different, based on Tukey's test for multiple comparisons ($p > 0.05$).
